# Children Who Develop Celiac Disease Exhibit Distinct Metabolic Pathways Among Their Gut Microbiota Years Before Diagnosis

**DOI:** 10.1101/2024.06.13.598929

**Authors:** Kristina Kelley, Dogus Dogru, Qian Huang, Yi Yang, Noah W. Palm, Johnny Ludvigsson, Emrah Altindis

## Abstract

Celiac disease (CD) is an autoimmune condition caused by a loss of tolerance to gluten in genetically predisposed individuals. While 30-40% of people possess the predisposing alleles, only 1-2% of the population is diagnosed with CD. This indicates environmental factors play a role in the pathogenesis of the disease, however the trigger of gluten tolerance loss is unknown. The gut microbiome composition of CD patients differs in comparison to their healthy counterparts; however, a causal link has not been established. In this study, we examined the alterations in the composition of the gut microbiota in a retrospective, longitudinal cohort of 10 children at age 1, matched for sex, human leukocyte antigen (HLA) genotype and breastfeeding duration. All samples were obtained from the pediatric donors prior to diagnosis (CD progressors). We used Ig-A sequencing combined with 16S sequencing for samples obtained at age 1. We also identified the functional metabolic pathways enriched in CD progressors compared to the healthy controls at ages 1, 2.5 (n=15-16) and 5 (n=9-13) using data from a similar study that we previously completed. Our findings demonstrate that CD progressors have ASV-level alterations in their gut microbiome as early as the first year of life, including the increased presence of some taxa that have been previously been reported to be enriched in CD. Using PICRUSt analysis, we also showed that inflammatory- and pathogenicity-related functions are enriched in CD progressors’ gut microbiome years before diagnosis. These pathways include glycine, serine and threonine metabolism, N-glycan biosynthesis and fatty acid biosynthesis and beta-lactam resistance, which are potentially contributing to chronic inflammation in CD. Overall, our results indicate distinct metabolic pathways enriched in the gut microbiome of CD progressors years before diagnosis. Understanding these pathways could advance our understanding of CD pathogenesis and its link to the gut microbiome.

## INTRODUCTION

Celiac Disease (CD) is an autoimmune disease triggered by the ingestion of gluten in genetically predisposed individuals. It is characterized by small intestinal inflammation that leads to villous atrophy however the exact mechanism(s) leading to disease development are not fully understood. CD affects 1-2% of the population worldwide, making it one of the most common lifelong disorders in people of all ages, and its prevalence and incidence continue to increase^1–3^. In addition to adult cases, there is a rising incidence of pediatric CD in the US and Europe^4,5^. In a local example in the USA (Denver, Colorado), the prevalence of CD by age 15 was 3.1%, which is two-fold higher than the estimation in the adult population^3^. 20-40% of the population possess human leukocyte antigen (HLA)-DQ2 or (HLA)-DQ8 haplotypes that bind and present gliadin, the gluten component acting as an antigen in CD. However, only 1-2% of individuals develop CD^6^. Previous research including discordant results for CD in identical twin studies^7^ ^8^, along with immigrant studies^9^ support the indication that environmental factors play a significant role in CD pathogenesis^9^. Currently, the only way to manage CD is strict adherence to a gluten free diet (GFD), but 20-40% of patients do not respond to the GFD and continue to have persistent or recurrent symptoms^10–12^. Additionally, a GFD is extremely difficult to adhere to and can negatively affect child development and quality of life^12–15^. Therefore, there is an urgent need to define environmental factors that play a role in CD onset and to understand the molecular mechanisms underlying CD etiology.

The gut microbiome is directly involved in the development and maintenance of the immune system and host immune responses^15^. Further, gut microbiota is associated with autoimmune diseases via direct and indirect interactions with innate and adaptive immune cells^16,17^. This interaction potentially results in loss of immune tolerance, chronic inflammation, and immune response against host tissues^18,19^. We and others previously reported altered microbial^20–27^ and metabolite composition^22,27–31^ in both infant and adult CD patients as well as in those at risk for developing the disease. Despite significant differences in the results of these studies on the patients, one common conclusion is that CD patients have an altered gut microbiome compared to healthy controls. This includes a decrease in some beneficial genus or species at the ASV level known to have anti-inflammatory properties, such as *Bifidobacteria*^24,26^. Some studies have also reported an increase in some bacterial species known to contribute to intestinal permeability, such as *Bacteroides* species.^21,24,32,33^. These findings suggest that gut microbiota may play an important role in CD pathogenesis; however, no causal link has been identified yet.

In our recent study^34^, we investigated the role of gut microbiota and microbial metabolites in CD onset by analyzing fecal and plasma samples from CD progressors and controls in the All Babies in Southeast Sweden (ABIS) cohort. . We observed that CD progressors have a distinct got microbiota composition at ages 2.5 and 5, years before diagnosis. We performed IgA sequencing and discovered that not only did CD progressors have more bacteria coated in IgA, but the bacterial was also differentially targeted by the immune system. Plasma metabolome analysis at age 5 identified 26 metabolites significantly altered in CD progressors, with the microbiota-derived metabolite taurodeoxycholic acid (TDCA) being the most significantly altered one in CD progressors. In vivo experiments showed that TDCA treatment induced villous atrophy and triggered an inflammatory immune response in vivo.

Here, we extended our analysis to include samples from the same cohort at age 1 to determine early microbiome alterations. We also used IgA-sequencing to determine the targets of IgA response in the gut microbiome. Furthermore, employing Phylogenetic Investigation of Communities by Reconstruction of Unobserved States (PICRUSt), we combined data from this and our previous study to identify microbial functions enriched in CD progressors at ages 1, 2.5, and 5. We demonstrated that although differences in gut microbiome composition are minimal, distinct metabolic pathways are enriched at age 1, while the opposite is observed for ages 2.5 and 5.

## METHODS

### Human Fecal and Plasma Samples

The fecal samples were obtained from subjects in the All Babies in Southeast Sweden (ABIS) cohort. ABIS study was ethically approved by the Research Ethics Committees of the Faculty of Health Science at Linköping University, Sweden (Ref. 1997/96287 and 2003/03-092) and the Medical Faculty of Lund University, Sweden (Dnr 99227, Dnr 99321). All children born in southeast Sweden between 1^st^ October 1997 and 1^st^ October 1999 were recruited. Informed consent from the parents was obtained. Fresh fecal samples were collected either at home or at the clinic. Samples collected at home were stored at -20 °C with freeze clamps, mailed to the WellBaby Clinic and stored dry at −80 °C. The questionnaire was completed by the parents to collect participants’ health information including, but not limited to, breast feeding duration, antibiotic use, gluten exposure time, and more. In total 10 fecal samples were collected for the analysis at age 1. "The small sample size is due to ABIS parents finding it difficult to collect stool samples at this age, as well as matching for HLA, breastfeeding, sex, and age with the controls.

### Study Cohort

All Babies in Southeast Sweden (ABIS) is a prospective population-based study that established a large biobank of biological specimens obtained longitudinally at birth and ages 1, 2.5, and 5. Previously, we reported on gut microbiome differences at ages 2.5 and 5^34^. In total, 17 000 children (78.6%) out of 21700 born in southeast Sweden Oct 1st 1997-1999 were included after their parents had given their informed consent. Among this large cohort, 230 children were diagnosed with celiac disease by the end of 2017, validated from the Swedish National Diagnosis Register (SNDR). CD diagnosis was made based on international classification of disease (ICD) codes-10 K90.0 according to the SNDR and met the ESPGHAN CD criteria but detailed information on height of TTG or any biopsy data with Mash stage was not available. To determine the role of gut microbiota in CD pathogenesis, we used ABIS samples selecting a sub cohort of 10 individuals, five of which developed CD but were not diagnosed with any other autoimmune disease as of December 2017. We selected these 10 subjects for age 1 because fecal samples obtained from these subjects at ages 2.5 and 5 were analyzed in our previous study (**Table S1**). The diagnosis of CD for the 5 subjects occurred after the last sample collection at ages 1.8, 6.4, 9.9, 12.6 and 13.2 (**Table S1**). Although we did not match subjects for other parameters, the timing of gluten exposure, delivery method, breastfeeding duration, family history of CD, infection history and use of antibiotics were comparable between groups (**Table S1**). The diagnosis of CD was confirmed at least twice in accordance with the Swedish National Diagnosis Register (https://doi.org/10.1186/1471-2458-11-450).

### IgA+ and IgA-Bacteria Separation

IgA-positive (IgA+) and IgA-negative (IgA-) bacteria were separated according to previously established methods onset^34^. In summary, frozen human fecal samples were placed in Fast Prep Lysing Matrix D with ceramic beads (MP Biomedicals) and incubated in 1 ml of Phosphate Buffered Saline (PBS) per 100 mg of samples on ice for 5 minutes to hydrate, followed by homogenization using bead beating for 7 seconds (Minibeadbeater; Biospec). The samples were then centrifuged at 50g for 10 minutes at 4°C to remove large debris. Fecal bacteria in the supernatants were collected (200 μl/sample) and washed three times with 500 μl PBS containing 1% (w/v) Bovine Serum Albumin (BSA, American Bioanalytical), then centrifuged for 5 minutes (6,000 x rpm, 4°C). A portion of this washed bacterial suspension (50 μl) was reserved as the pre-sorting sample for 16S sequencing analysis. Following washing, bacterial pellets were re-suspended in 50 μl of blocking buffer (PBS containing 1% (w/v) BSA and 20% Normal Mouse Serum, Jackson ImmunoResearch), and then incubated for 20 minutes on ice.

Subsequently, they were stained with 100 μl of PE-conjugated mouse anti-human IgA (1:40; Miltenyi Biotec clone IS11-8E10) for 30 minutes on ice. After staining, the samples underwent three washes with 500 μl of BSA solution containing 1% (w/v) before proceeding to either flow cytometry analysis or cell separation.. PE anti-human IgA stained bacteria were incubated with Anti-PE Magnetic Activated Cell Sorting (MACS) beads (Miltenyi Biotec) (1:5) for 30 minutes on ice and then separated by a custom magnetic plate for 10 minutes on ice. Fecal bacteria bound to the magnetic plate were collected as IgA+ samples for 16S sequencing analysis. Stained and MACS bead-bound bacteria unbound to the magnet were collected (20∼40 μl) and passed through MACS molecular columns (Miltenyi Biotec) (one sample/column), followed by flushing with 480 μl PBS containing 1% (w/v) BSA. The total pass-through (∼500 μl) was loaded onto columns once more. The columns were flushed with 500 μl PBS containing 1% (w/v) BSA. The total column pass-through (∼1 ml) was saved as IgA-samples for 16S sequencing analysis.

### Fecal IgA Flow Cytometry Analysis

Bacterial cells were extracted from fecal samples following the procedure outlined in the IgA+ and IgA-Bacteria Separation section of this manuscript. These cells were then subjected to staining with PE Anti-human IgA antibodies (1:100; Miltenyi Biotec clone IS11-8E10) for 30 minutes on ice. Following two washes, the bacteria were stained with TO-PRO®-3 (ThermoFisher Scientific) to distinguish them from fecal debris or particles. Subsequently, the stained bacteria were analyzed using a BD FACSAriaTM IIIu cell sorter (Becton-Dickinson) according to previously described methods^47^, categorizing them as TO-PRO®-3+IgA+/- cells.

### 16S rRNA Gene Sequencing

16S rRNA sequencing targeting the V4 region was conducted for all bacterial samples using the MiSeq platform, employing barcoded primers, as outlined in previous literature^48^. To summarize, bacterial samples were initially suspended in 90 μls of MicroBead Lysis Solution supplemented with 10% RNAse-A, then sonicated in a water bath at 50°C for 5 minutes. Subsequently, the samples were transferred to a plate containing 50 μls of Lysing Matrix B (MP Biomedicals) and homogenized via bead-beating for 5 minutes. Following centrifugation at 4122 x g and 4°C for 6 minutes, the supernatant was carefully transferred to 2 ml deep-well plates (Axygen Scientific). Bacterial DNA from the samples was extracted and purified using the MagAttract Microbial kit (QIAGEN), following the manufacturer’s instructions. PCR amplification of the V4 region of 16S ribosomal RNA was carried out in duplicate (3 μl purified DNA per reaction) with 33 cycles, utilizing Phusion DNA polymerase (New England Bioscience)^48^.

Subsequently, the amplified PCR products were normalized using the SequalPrepTM normalization plate kit (ThermoFisher Scientific) and then pooled. The concentration of the pooled library was determined using the NGS Library Quantification Complete kit (Roche 07960204001) before being loaded onto a MiSeq sequencer. Illumina MiSeq Reagent Kit V2 (500 cycles) was employed to generate 2x250bp paired-end reads. The raw reads were demultiplexed using Qiime1 (version 1.9), resulting in an average of 30,471 reads per sample.

### Bioinformatic Analysis and Statistics

Microbial diversity and statistical analyses involved the initial step of filtering and trimming the bacterial 16S rRNA amplicon sequencing reads. Subsequently, sample inference was conducted to convert the amplicon sequences into an Amplicon Sequence Variant (ASV) table using dada2, utilizing the Ribosomal Database Project Training Set 16^35^. Exploratory and inferential analyses were performed by using phyloseq^36^ and vegan^37^, which includes Non-metric MultiDimenstional Scaling (NMDS) analysis using Bray–Curtis dissimilarity, Principle Components Analysis (PCA), alpha and beta diversity estimates, and taxa agglomeration.

Differential OTU abundance was assessed per time point by edgeR^38^ with two-sided empirical Bayes quasi-likelihood F-tests tests. P-values were corrected by using the Benjamini-Hochberg false discovery rate (FDR), and FDR < 0.25 was considered statistically significant^39^. The prediction of gene content and pathway abundance were performed using the Kyoto Encyclopedia of Genes and Genomes (KEGG) database and PICRUSt2^40–42^. Differential KEGG pathway abundance was assessed by using limma^43^. The bar plots and box plots were made by using ggplot2^44^, and heatmap by pheatmap^45^.

### Data Availability

The 16S raw sequencing data generated in this study is available at NCBI Sequence Read Archive Bioproject PRJNA631001. The gut microbiome analysis codes generated in this study is available at this link: https://github.com/jdreyf/celiac-gut-microbiome.

## RESULTS

### Firmicutes are elevated in the gut microbiome of CD progressors age 1

In total, we identified 120 amplicon sequencing variants (ASVs) at age 1 (**Table S2**). While, there was a trend of increase in alpha diversity in the CD progressor group (observed ASV: unpaired t-test p=0.0887; Simpson index: unpaired t-test p=0.0787), consistent with other studies^46^, it was not significant (**Figure 1A**, upper panel). Similarly, beta diversity was also comparable (p=0.915) (**Figure 1A**, lower panel). Principal component analysis (PCA) plots showed a trend of separation of gut microbiome composition (**Figure 1B**). Relative abundance analysis revealed that CD progressors had higher levels of Firmicutes than controls (mean average abundant (MAA): CD=0.619, Ctrl=0.427; p=0.0148) (**Figure 1C**, upper panel, **Table S3**). On the other hand, there was no difference in other phylogenetic levels including at the genus level (**Figure 1C**, lower panel). Focusing on ASV-level differences, we found only 14 ASVs that differed at age 1 (**Figure 1E, Table S2**), with the majority remaining unchanged (FDR <0.1, p<0.05; **Figure 1D**). For example, *Ruminococcus bromii (*FC=3410, FDR=0.05), *Dialister invisus* (FC=1410, FDR=0.05) and *Bifidobacterium dentium* (FC=1060, FDR=0.05) were enriched in CD progressors along with some ASVs from the *Clostridium* genus including Clostridium XVII (FC=4560, FDR=.05), and *Clostridium XIV scindens* (FC=309, FDR=.05 ). ). interestingly, most of these *Clostridium* species are related to TDCA production^34^ that we recently identified a potential role in CD onset^34^ . Conversely, Enterococcus (FC=-1000, FDR=0.05) was enriched in control samples (**Figure S1, Figure 1D & E**). However, it’s noteworthy that there was considerable variability in the occurrence of certain ASVs within both groups.

**Figure 1.**
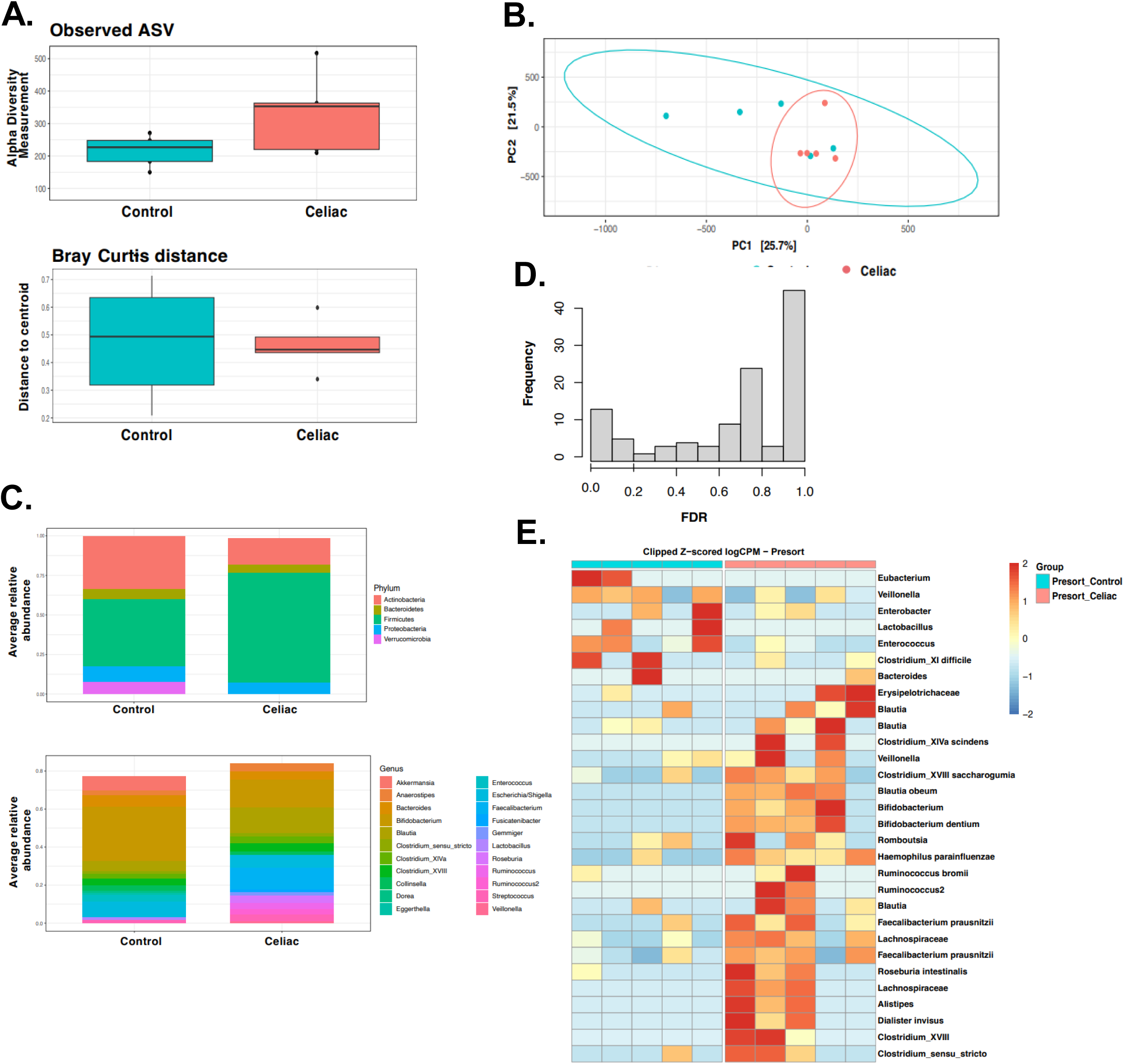
The gut microbiota of CD progressors is distinct at the ASV level. **A.** Box plots showing the comparison between CD progressors (n=5) and healthy controls (n=5) the alpha diversity measured by observed ASVs (upper panel) and the beta diversity measured by Bray–Curtis dissimilarity (lower panel). Statistical analysis was performed using ANOVA (alpha diversity) and PERMAMOVA (Bray-Curtis distance). **B.** Principal component analysis (PCA) ordination of sample similarity/dissimilarity between CD progressors and healthy controls at age 1 year. Each circle represents an individual CD progressor (red) or control (blue) sample. **C.** Average relative abundance of bacterial phylum (upper panel) or genera (lower panel) of greater than 1% abundance (proportion) between the gut microbiota of CD progressors and healthy controls. (taxa average relative abundance>1%). Statistical analysis was performed using two-tailed *t* tests with the Benjamini and Hochberg method to control the false discovery rate (FDR). **D.** Empirical Bayes quasi-likelihood F-tests analysis for the comparisons of gut microbiota ASVs between CD progressors and healthy controls . Frequency: number of ASVs. **E.** Heat map showing the relative abundance of the top ASVs differentially enriched in CD progressors and healthy controls. Each column represents an individual sample and each row represents an ASV.

### CD progressors have a different IgA response at age 1

To identify microbiota species highly coated with IgA and determine whether there is any difference in the IgA response at age 1, we used a modified method of IgA-sequencing^34^. In the IgA- group, the alpha diversity was higher (observed ASV: unpaired t-test p= 0.0121; Simpson index: unpaired t-test p= 0.0828) for the CD progressors compared to the controls (**Figure 2A**, upper panel), while the alpha diversity of the IgA+ fractions were similar (**Figure 2A**, upper panel). Beta diversity was comparable in all groups (**Figure 2A**, lower panel). PCA analysis showed a trend of separation between IgA+ and IgA-bacteria both in control and CD samples (**Figure 2B).** IgA+ bacteria accounted for 4.57% (LS mean) at age 1 in controls (**Figure 2C**) and 10.99% at age 1 in CD progressors, though this difference was not statistically significant (p=0.24). Consistent with the pre-sorting data, CD progressors had higher levels of Firmicutes than controls for both IgA+ and IgA-fractions (for IgA+ MAA: CD= 0.4702, Ctrl= 0.4097; p=0.0422 and for IgA-MAA: CD= 0.6898; Ctrl= 0.4097; p= 0.0171) (**Figure 2D**, upper panel, **Table S2**) and we did not observe any differences at other phylogenetic levels (**Figure 2D**, lower panel).

**Figure 2.**
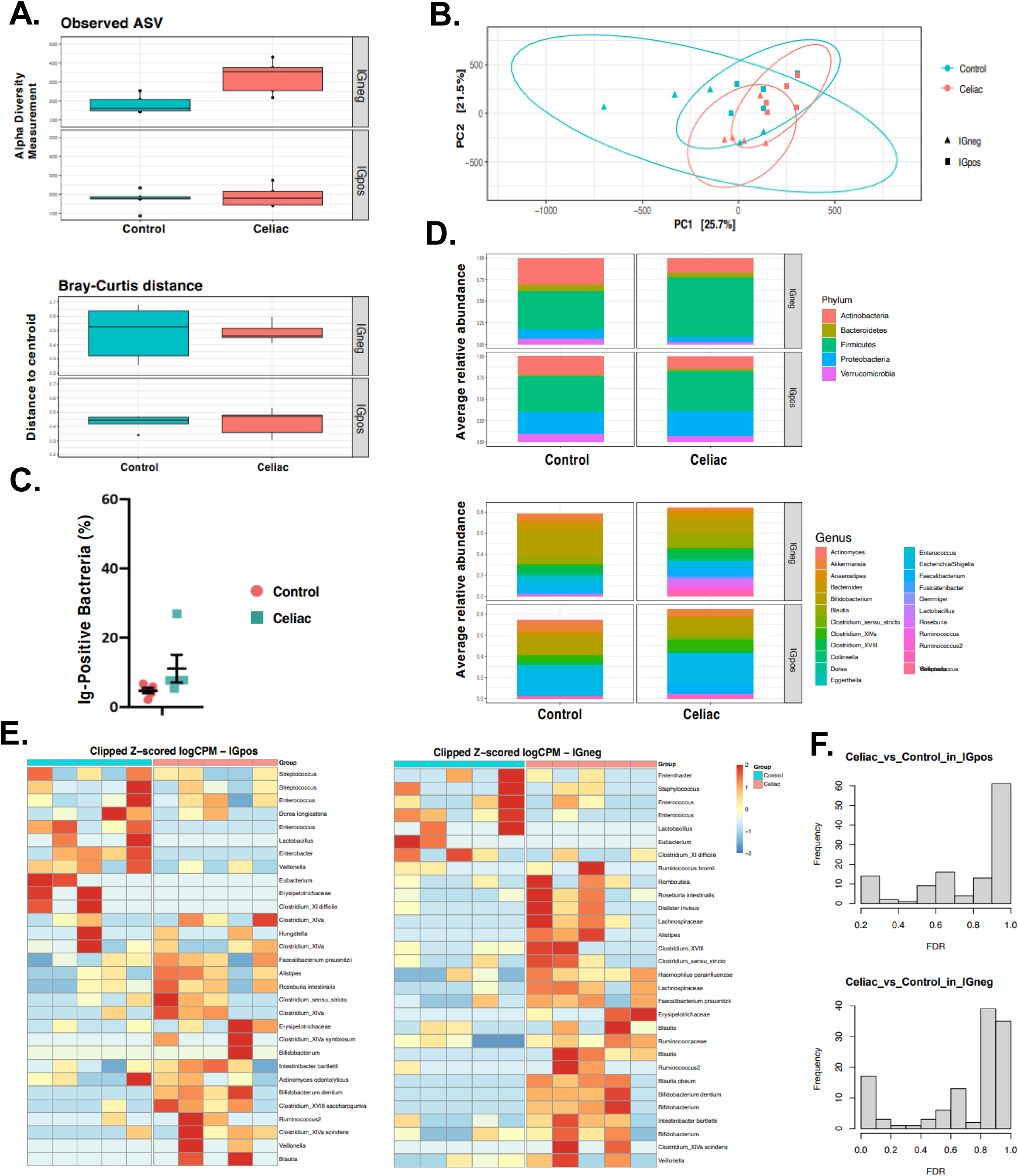
IgA-based sorting and 16S sequencing revealed that gut microbiota were differentially coated by IgA. **A.** Box plots showing the comparison between CD progressors and healthy controls: the alpha diversity measured by observed for IgA+/IgA- microbiota (upper panel), and the beta diversity measured by Bray–Curtis dissimilarity for IgA+/IgA- microbiota (lower panel) at age 1 for CD progressors (red) and healthy controls (blue). Statistical analysis was performed using ANOVA (alpha diversity) and PERMAMOVA (Bray-Curtis distance). **B.** Principal component analysis (PCA) of sample similarity/dissimilarity between IgA+ (square) and IgA- (triangle) microbiota in healthy control (blue) or CD progressors (red). Each point represents an individual sample. **C.** Percentage of IgA positive bacteria recovered ( blue) and healthy controls (red) . Indicated are mean ±SEM. Statistical analysis was performed using t wo way ANOVA **D.** Average relative abundance of IgA+/IgA- bacterial phylum (upper) or genera (lower) of greater than 1% abundance (proportion) between the gut microbiota of CD progressors and healthy controls (taxa average relative abundance>1%). Statistical analysis was performed using two-tailed t-tests with Benjamini and Hochberg method to control False Discovery Rate (FDR). **E.** Heat map showing the relative abundance of the top ASVs significantly different between IgA+ (left) IgA- (right) CD progressors (red) and healthy controls (blue) . Each column represents an individual sample and each raw represents an ASV. **F.** Empirical Bayes quasi-likelihood F-tests analysis for the comparisons of IgA-coated (top panel) or non-coated (bottom panel) gut microbiota ASVs between CD progressors and healthy controls.

Notably, we did not identify any differences between IgA- and IgA+ fractions of controls or CD progressors. There was also no difference between the IgA+ fractions of both groups. However, several taxa were differentially abundant (FDR<0.1) between CD progressors and healthy controls in the IgA-negative microbiota (**Figure 2E** and **2F**) following our observations in the basal microbiome differences. Erysipelotrichacea (FC=2170, p=0.00438, FDR=0.0739), Ruminococcus2 (FC=1800, p= 0.0103, FDR= 0.0775), *B. dentium* (FC=1720, p=0.00332, FDR=0.0739), *Roseburia intestinalis* (FC=1670, p=0.00517, FDR=0.0739), and Clostridium_XVIII (FC=1310, p=0.00888, FDR=0.0761), were enriched in the IgA- microbiota of CD progressors compared to healthy controls. In contrast, three taxa were decreased in the IgA- gut microbiota of CD progressors including Enterococcus (FC=-2900, p=0.00323, FDR=0.0739), Lactobacillus (FC=-1410, p=0.0069, FDR=0.0739), and Eubacterium (FC=-809, p=0.00606, FDR=0.0739) (**Table S2**) .

### Pathogenesis and Inflammation Related Functions Are Enriched in CD Progressors’ Gut Microbiota

PICRUSt analysis^47^ is designed to estimate the functional metagenome of gut bacteria using 16S rRNA data. In this study, we utilized PICRUSt to examine enriched functional pathways in CD progressors. To this end, we combined the sequencing data from CD progressors at age 1 in this study and ages 2.5, and 5 reported in our previous study^34^,noting that all sample preparation and sequencing were conducted simultaneously for consistent results. Combining PICRUSt with Kyoto Encyclopedia of Genes and Genomes (KEGG) metabolic pathway analysis, we identified 71 different metabolic pathways that differed between CD and control samples at age 1 (FDR<0.05, **Figure 3A, Table S5**). Among these pathways, N-glycan biosynthesis (FC=3.42, FDR=0.0208), penicillin and cephalosporin biosynthesis (FC=2.75, FDR=0.0259), beta-Lactam resistance (FC=2.51, FDR=0.0208) and bacterial chemotaxis (FC=2.28, FDR=6.47e-4) pathways were among the top pathways enriched in CD progressors. Interestingly, most of these pathways are involved in bacterial pathogenesis^48,49^ or shaping the composition of microbiota^50,51^. Additional pathways that were highly enriched at age one included the fatty acid biosynthesis pathway (FC=1.92, p= 0.000217, FDR=0.0082), pathways related to D-glutamine and D-glutamate metabolism (FC=1.66, p= 0.00397, FDR=0.0208), nitrogen metabolism (FC=1.6, p= 0.0354, FDR=0.0082), glyoxylate and dicarboxylate metabolism (FC=1.92, p= 0.00513, FDR=0.0209), glycine, serine, and threonine metabolism (FC=1.51 p=0.0122, FDR=0.0296). We also identified 9 pathways showing trends of difference both at age 2.5 and at age 5 (**Table S5)**. For example, styrene degradation, lysine degradation, fatty acid metabolism and glutathione metabolism were decreased in CD progressors at age 2.5 (P<0.05). Meanwhile, retinol metabolism, steroid hormone biosynthesis, and glycosaminoglycan degradation pathways were increased in CD progressors at age 5 (P<0.05).

**Figure 3.**
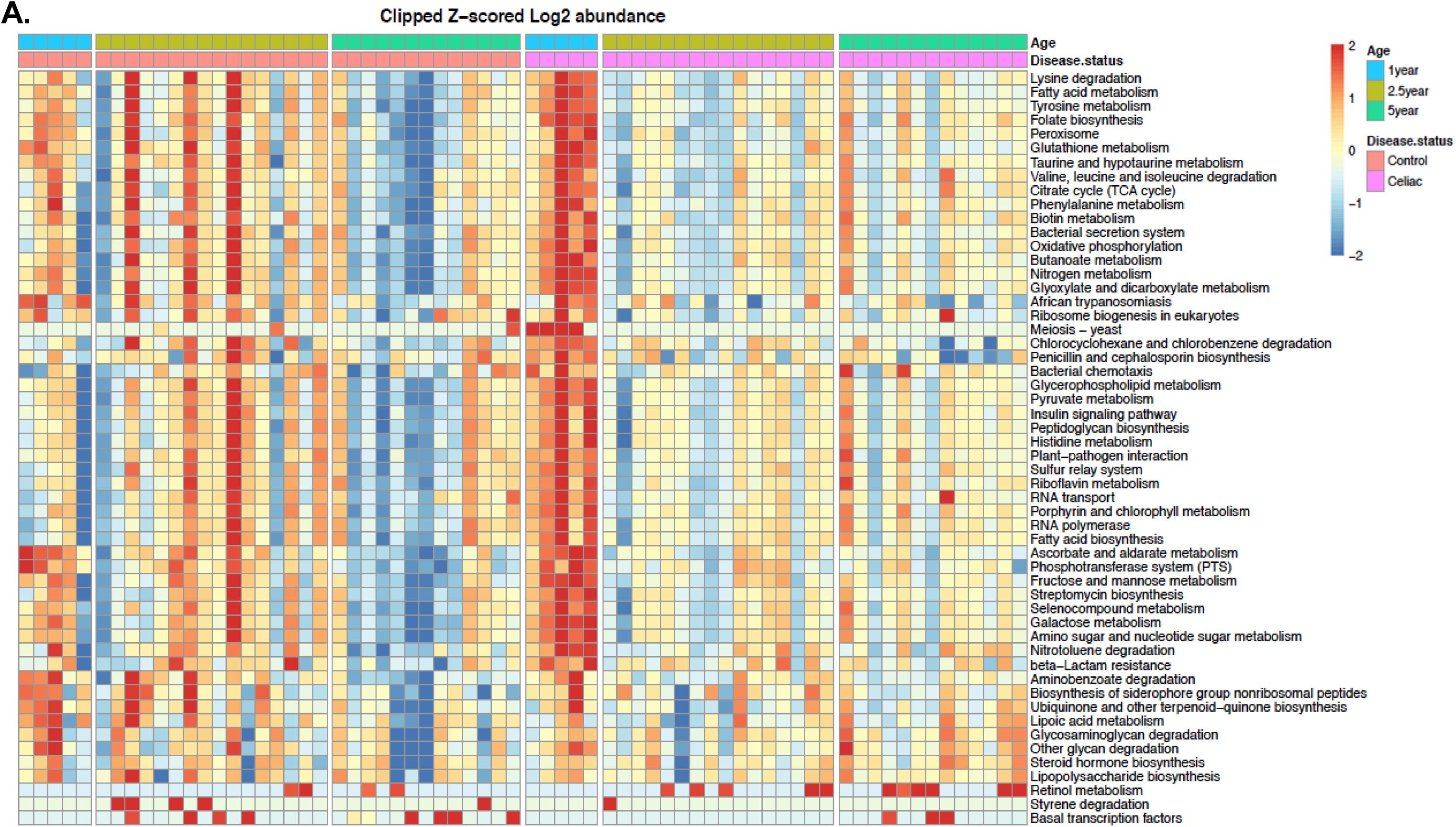
The enriched microbial pathways altered in CD progressors’ gut microbiota are significantly different from healthy controls. **A.** Heat map of PICRUSt predicted metabolic pathways of CD progressors and healthy controls. Each column represents an individual participant and each row represents a predicted microbial functional pathway. Color code is shown on the figure. Ages 1year old : n=5/group); age 2.5 years old: n=16/group; age 5 years old: n=13/group.

We also used PICRUSt to examine the functional pathways comparing IgA+ to IgA- microbiota (**Figure 4A, S3, Table S5)**. We identified styrene degradation pathway enriched in IgA+ population at age 1. Further, we identified 31 different functional pathways at age 2.5 and 23 functional pathways at age 5 (FDR<0.05) in the healthy subjects. The top enriched pathways in healthy IgA+ population were styrene degradation (age 2.5: FC=103, FDR=1.97e-12; age 5: FC=77.1, FDR=4.92e-9), chloroalkane and chloroalkene degradation (age 2.5: FC=534, FDR=9.09e-11; age 5: FC=222, FDR=1.42e-6), and toluene degradation at the ages (age 2.5: FC=482, FDR=2.3e-7; age 5: FC=99.6, FDR=1.73e-3). Analyzing CD samples, we identified 7 pathways at age 1, 28 pathways at age 2.5 and 13 pathways at age 5 enriched in IgA+ microbiota (FDR<0.05). The most significantly enriched pathways in CD progressors’ IgA+ microbiota were beta-alanine metabolism (age 1: FC=1210, FDR=0.033; age 2.5: FC=145, FDR=7.43e-4; age 5: FDR>0.05), chloroalkane and chloroalkene degradation (age 1: FC=997, FDR=4.86e-5, age 2.5: FC=148, FDR=1.83e-7; age 5: FC=714, FDR=2.44e-8), and styrene degradation (age 1: FC=856, FDR=1.87e-8, age 2.5: FC=421, FDR=1.42e-19; age 5: FC=440, FDR=8.43e-15). At age 1, CD progressors had more metabolic pathways predicted to be enriched in the IgA+ samples compared to healthy control. For example, pathogenic pathways, bacterial invasion of epithelial cells (FDR=0.0174, FC=16.2) and beta-alanine metabolism (FDR=0.033, FC=1210) were identified in CD IgA+ microbiota population but was absent in control IgA+ at age 1.

**Figure 4.**
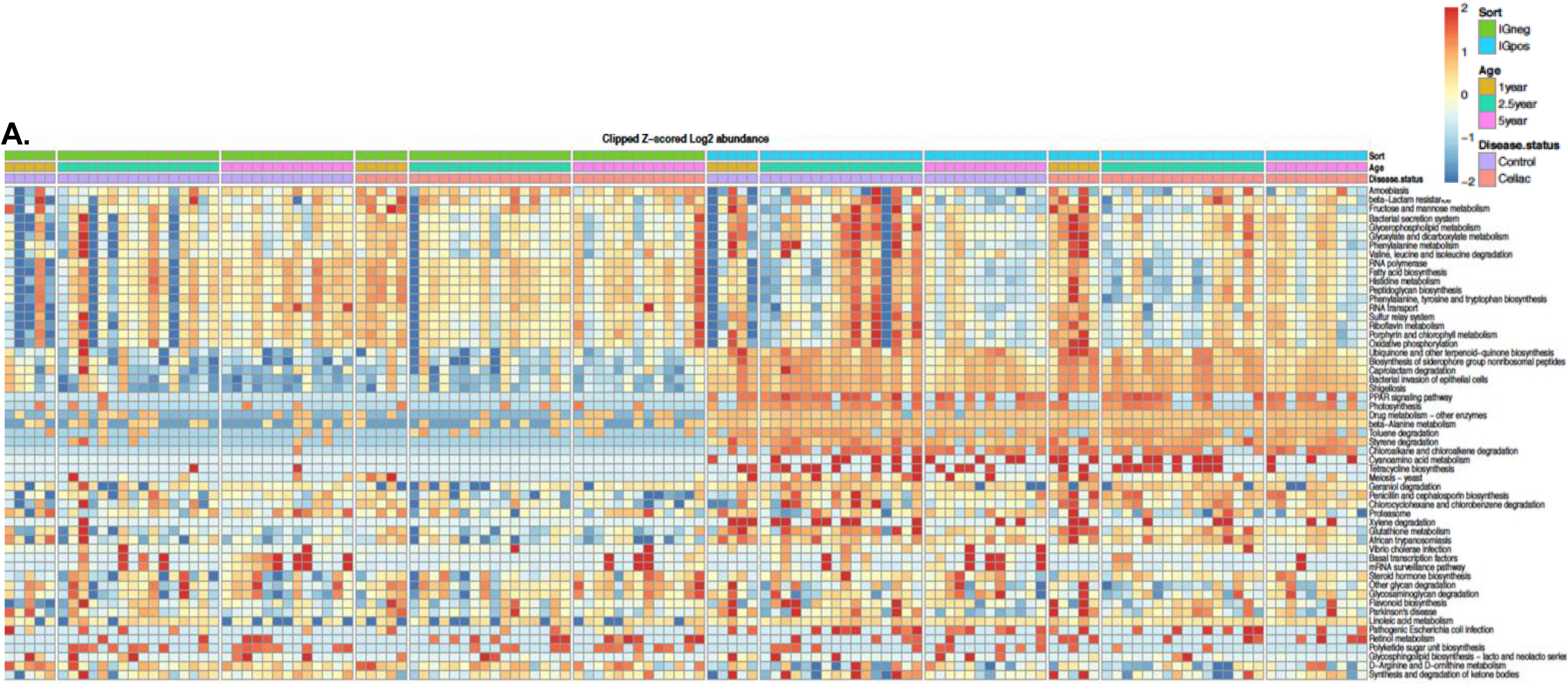
The enriched microbial pathways altered in CD progressors’ gut microbiota are significantly different between IgA+ and IgA- bacteria. **A.** Heat map of predicted metabolic pathways of CD progressors and healthy controls obtained from PICRUSt analysis after IgA sequencing. Each column represents an individual participant and each raw represents a predicted microbial functional pathway. Color Code is shown on the figure. Ages 1year old : n=5/group); age 2.5 years old: n=16/group; age 5 years old: n=13/group.

## DISCUSSION

Recent studies have demonstrated strong associations between the gut microbiota and the pathogenesis of autoimmune diseases including Type 1 diabetes^56,57^, multiple sclerosis^58,59^, and rheumatoid arthritis^60,61^. Studies of the gut microbiome in CD have demonstrated intestinal dysbiosis in CD patients^20,22–27,62,63^. However, most of these studies utilized adult samples from individuals already diagnosed with the disease rather than adopting a longitudinal, prospective approach with samples collected prior to disease onset, as this study. Human gut microbiota development is divided into three phases; a developmental phase (months 3-14), a transitional phase (months 15-30), and a stable phase (months 31-46)^64^. Because recent data show that many childhood CD cases will develop in the first years of life,^65^, we analyzed samples representing all of these critical phases. We previously reported that CD progressors at age 2.5 and 5 have a distinct gut microbiome composition and differential IgA response, and at age 5, have an altered cytokine profile and plasma metabolome. Here, we report, for what we believe is the first time, that there are significant differences in the predicted functional levels at the developmental phase in CD progressors’ gut microbiome. Consistent with previous reports^63^ , we identified the proportion of phylum Firmicutes higher in CD progressors at age 1 (**Figure 1C**). Bacterial proteases of species mostly classified within the Firmicutes phylum are involved in gluten metabolism and this might be a potential link to CD^66^. Additionally, we identified bacterial species enriched in CD progressors at age 1 including *Bifidobacterium dentium*, *Clostridium XIVa sciendens*, *Faecalbacterium prausnitzii, Lachnospiraceae,* and *Alistipes,* and a trend of increase in *Clostridium sensu stricto,* however there was variation between samples. This is consistent with previous research findings that show an increase of *Faecalbacterium prausnitzii* in untreated CD^67^, as well as *Lachnospiraceae* in CD progressors^30^, and infants at high risk of developing the disease^68^. Additionally, *Alistipes* was increased in CD patients^67,69^ . Notably, a previous study reported that the abundance of *R. bromii* was greatly reduced in CD patients when gluten free (GF) diet was introduced^70^. Further, infants at high risk for developing CD had higher levels of *Clostridium sensu stricto*^71^. We found that several taxa were decreased in the age 1 CD progressor samples, such as Veillonella, Enterobacteriaceae, and Bacteriodes. This is consistent with other studies showing Veillonella decreased in CD progressors^72^ , Enteriobacter decreased in infants at high risk for CD^71^ and Bacteroides decreased in both CD progressors ^72^ and patients^69^. Consistent with our findings, other studies have shown that the abundance of *B. dentium* was increased in the CD patients^73^. *Clostridium XIVa* genera is responsible for producing the proinflammatory metabolite TDCA, which might be implicated in CD pathogenesis. In our previous study, we showed that TDCA was enriched in serum of CD progressors at age 5. Additionally, treating B6 mice with TDCA caused an altered immune response and villous atrophy that is characteristic of CD^34^. In this study, we showed that several Clostridium members were enriched in CD progressors. When we compared our data in this study (age 1) to the data obtained from our previous study (ages 2.5 and 5), we found that the differences in the gut microbiota were less drastic at the phylum level, but significantly increased in the ASV level by age.

Intestinal IgA plays a crucial role in defending against pathogenic microorganisms and in maintaining gut microbiome homeostasis. Interestingly, IgA-deficient patients are more susceptible to variety of pathologies, including CD^75^. Planer et al described mucosal IgA responses progression during two postnatal years in healthy US twins^76^. They showed that (i) IgA coated bacteria is affected by age and host genetics and (ii) IgA response is determined by "intrinsic" properties of gut microbiota community members. We used a similar approach to investigate the gut immune development towards healthy and CD states, we initially focused on the development of the IgA response during the gut microbiota maturation. At age 1, we did not identify any ASVs that were significantly different in IgA- and IgA+ samples. While it is potentially related to small sample size used in this study, it might also suggest that the intestinal IgA response is not mature enough to target specific bacteria in the gut. However, we showed that the IgA response is highly selective and only a small fraction of the gut microbiota is highly coated with IgA in the first year of life. This is consistent with our previous findings at age 2.5 and 5 and notably, we found a significant increase in the percentage of IgA coated bacteria in CD progressors at age 5^34^. While a reduction of secretory IgA (sIgA) using infant (4-6 months) fecal samples in CD progressors^27^ was reported previously, we did not identify any significant difference in the ratio of IgA coated bacteria at age 1.

The abundance analysis (**Table S2, Fig 2E**) showed that CD progressors’ IgA+ microbiota is enriched with Firmicutes. Consistent with this finding, previous studies reported higher proportions of Firmicutes in HLA high-risk infants^29,71^. Notably, most of the bacterial strains in the human gut microbiota that can metabolize gluten are classified with the Firmicutes, which includes strains that have been found to show extracellular proteolytic activity against gluten proteins^72^. Bacterial metabolism of gluten might result in different gluten peptide fragments, potentially related to CD autoimmunity.

PICRUSt analyses showed significant differences at all developmental phases, in particular, within the transition period at age 1. Most significant differences were identified in pathways related to bacterial pathogenesis and shaping the composition of microbiota. For example, glutathione metabolism was greatly decreased in CD progressors. Decreased glutathione redox cycle in CD patients is strongly associated with disease development^77^. Several other pathways found enriched in age 1 samples have previously been found to be upregulated in CD patients^55^, including pathways related to D-glutamine and D-glutamate metabolism, nitrogen metabolism, glyoxylate and dicarboxylate metabolism, glycine, serine, and threonine metabolism. An over production of nitric oxide (NO), a product of nitrogen metabolism, has been associated with the disruption of gut microbiota and directly linked with autoimmune conditions involving gut inflammation, such as Crohn’s disease and ulcerative colitis^78,79^. The fatty acid biosynthesis pathway was significantly increased at age 1. Alterations in the levels of fatty acids is associated with gastrointestinal diseases such as CD^80^. Fatty acids can modulate function of immune cells and inflammatory processes both directly and indirectly^81^. In particular, short chain fatty acids (SCFAs) are metabolites produced by the gut microbiota and play a role in gut barrier maintenance. The alteration and accumulation of some SCFAs may be associated with untreated CD^82^. At age 5, PICRUSt predicted a trend that retinol metabolism, steroid hormone biosynthesis, and glycosaminoglycan degradation as over-represented pathways in CD progressors. Retinoic acid is one of the products of retinol metabolism and plays a key role in the intestinal inflammatory response^83^ as well as impacting the expression of cytokine expression^84^. A previous study showed that retinoic acid mediated inflammatory responses to gluten in both *in vitro* and *in vivo* studies^85^. The increased retinol metabolism and glycosaminoglycan degradation pathways in CD progressors are potentially related to chronic inflammation. Indeed, glycosaminoglycan helps to form a protective barrier for the intestinal mucin. The breakdown of glycosaminoglycan is reported to be associated with inflammatory response in intestinal disorders such as IBD^86^.

We also used PICRUSt after sorting via IgA sequencing to compare the functional pathways associated with the IgA+ and IgA- bacteria. Among our CD progressor samples, we identified 7 pathways at age 1, 28 pathways at age 2.5 and 13 pathways at age 5 enriched in IgA+ microbiota. Among these were a pathway involved in the bacterial invasion of epithelial cells, which was predicted in the IgA+ microbiota, but not the IgA- microbiota at age 1.

The gut microbiome develops in the first years of life by progressing through several distinct phases. Entering the transition phase, the gut microbiota in CD progressors displayed more proinflammatory and oxidative stress related features. At stable phases, gut microbiota in CD progressors begins to become more involved in functions related to the clinical manifestation of the disease, such as those associated with intestinal inflammation. This longitudinal observation provides insight into the proinflammatory and pathogenic function of gut microbiota in different stages of early CD pathogenesis.

Currently, the only way to treat CD is strict adherence to a gluten-free (GF) diet, but 20% of patients do not respond to GF diet and continue to have persistent or recurrent symptoms^10^ . CD permanently reshapes intestinal immunity and alterations in TCRγδ+ intraepithelial lymphocytes in particular may underlie non-responsiveness to the GF diet^87^. Our findings suggest that enriched pathways identified in the CD progressors’ gut microbiota might be related or contribute to an inflammatory environment that could be an important component of intestinal inflammation in CD. The proinflammatory pathways identified in this study could potentially trigger local and systemic inflammation independent of the diet and may explain a failure to respond to GF diet in some patients.

The main strength of this study lies in the longitudinal sampling that represents all three phases of gut microbiota development in children for the PICRUST analysis. Further, applying IgA-seq analysis, for what we believe is the first time, we examined an important dimension to evaluate CD pathogenesis. The main limitation of this exploratory study is the small sample size in the gut microbiome cohort, particularly at age 1 (n=5). Using 16S-sequencing is another limitation that requires PICRUST to predict functional pathways instead of shotgun sequencing directly acquiring the gene information. Therefore, follow up studies with large sample size and deep shotgun sequencing are needed to validate our findings, increasing the statistical power and depth. Taken together, our findings suggest that the gut microbiota of CD progressors in the first five years of life has functionally different gut microbiota and this can potentially contribute to the onset and progression of CD. We identified a small number of ASVs at age 1 that were differentially present in the CD progressors microbiota, some of which have been previously reported to be associated with those at high risk for developing CD or patients in active CD. We also identified several functional pathways enriched in CD progressors at each stage of microbiome development that are involved in bacterial pathogenesis, shaping the microbiota composition, and gut inflammation, as well as some that have been previously linked to CD. Understanding the role of the gut microbiota in chronic inflammation in CD and targeting inflammatory bacteria or developing anti-inflammatory probiotics/prebiotics may open novel approaches to understand disease pathogenesis and reveal new preventive and treatment models.

## Acknowledgements

We are grateful to all children participating in the ABIS study, and their parents. Thanks also to Ingela Johansson, KEF, Linköping, for her skillful work with the samples, and Åshild Faresjö for register data. The authors also want to thank Hui Pan, Jonathan Dreyfuss (Joslin Diabetes Center Bioinformatic Core) for their help with bioinformatic analysis and statistics. The authors would also like to acknowledge Patrick Autissier for the cytometry service (Flow Cytometry Core of Boston College). This work was supported by a G. Harold & Leila Y. Mathers Foundation grants to EA. The ABIS-study has been supported by Swedish Research Council (K2005-72X-11242-11A and K2008-69X-20826-01-4) and the Swedish Child Diabetes Foundation (Barndiabetesfonden), JDRF Wallenberg Foundation (K 98-99D-12813-01A), Medical Research Council of Southeast Sweden (FORSS) and the Swedish Council for Working Life and Social Research (FAS2004–1775) and Östgöta Brandstodsbolag.

## Author contributions

E.A. designed the research. K.K. and E.A. wrote the paper. Q.H., Y.Y., and N.W.P. assisted with 16S-sequencing and IgA-seq experiments and analysis. K.K. assisted with data analysis. D.D. contributed to the bioinformatics analysis. J.L. is the Head of the ABIS study and assisted with human fecal sample collection, classification, and data interpretation. All authors contributed to data analysis, approved the final version of the manuscript, and participated in producing it.

## Supplementary Figure Legends

**Figure S1. Violin plot representation of most enriched genus/species in CD progressors compared to healthy controls.** Violin plots showing the abundance of ASVs in CD progressors (n=5) and healthy controls (n=5) with False Discovery Rate less than 0.1. Statistical significance was determined using empirical Bayes quasi-likelihood F-tests.

## REFERENCES

1 Almallouhi, E. et al. Increasing Incidence and Altered Presentation in a Population-based Study of Pediatric Celiac Disease in North America. J Pediatr Gastroenterol Nutr 65, 432–437, doi:10.1097/MPG.0000000000001532 (2017).

2 Mariné, M. et al. The prevalence of coeliac disease is significantly higher in children compared with adults. Aliment Pharm Ther 33, 477–486, doi:10.1111/j.1365-2036.2010.04543.x (2011).

3 Liu, E. et al. High Incidence of Celiac Disease in a Long-term Study of Adolescents With Susceptibility Genotypes. Gastroenterology 152, 1329–1336 e1321, doi:10.1053/j.gastro.2017.02.002 (2017).

4 King, J. A. et al. Incidence of Celiac Disease Is Increasing Over Time: A Systematic Review and Meta-analysis. Am J Gastroenterol 115, 507–525, doi:10.14309/ajg.0000000000000523 (2020).

5 Myleus, A. et al. Celiac disease revealed in 3% of Swedish 12-year-olds born during an epidemic. J Pediatr Gastroenterol Nutr 49, 170–176, doi:10.1097/MPG.0b013e31818c52cc (2009).

6 Wolters, V. M. & Wijmenga, C. Genetic background of celiac disease and its clinical implications. Am J Gastroenterol 103, 190–195, doi:10.1111/j.1572-0241.2007.01471.x (2008).

7 Greco, L. et al. The first large population based twin study of coeliac disease. Gut 50, 624–628, doi:10.1136/gut.50.5.624 (2002).

8 Kuja-Halkola, R. et al. Heritability of non-HLA genetics in coeliac disease: a population-based study in 107 000 twins. Gut 65, 1793–1798, doi:10.1136/gutjnl-2016-311713 (2016).

9 Cataldo, F. et al. Epidemiological and clinical features in immigrant children with coeliac disease: an Italian multicentre study. Dig Liver Dis 36, 722–729, doi:10.1016/j.dld.2004.03.021 (2004).

10 Stasi, E. et al. Frequency and Cause of Persistent Symptoms in Celiac Disease Patients on a Long-term Gluten-free Diet. J Clin Gastroenterol 50, 239–243, doi:10.1097/MCG.0000000000000392 (2016).

11 Veeraraghavan, G. et al. Non-responsive celiac disease in children on a gluten free diet. World J Gastroenterol 27, 1311–1320, doi:10.3748/wjg.v27.i13.1311 (2021).

12 Roos, S., Liedberg, G. M., Hellstrom, I. & Wilhelmsson, S. Persistent Symptoms in People With Celiac Disease Despite Gluten-Free Diet: A Concern? Gastroenterol Nurs 42, 496–503, doi:10.1097/SGA.0000000000000377 (2019).

13 West, J., Logan, R. F., Card, T. R., Smith, C. & Hubbard, R. Risk of vascular disease in adults with diagnosed coeliac disease: a population-based study. Aliment Pharmacol Ther 20, 73–79, doi:10.1111/j.1365-2036.2004.02008.x (2004).

14 Midhagen, G. & Hallert, C. High rate of gastrointestinal symptoms in celiac patients living on a gluten-free diet: controlled study. Am J Gastroenterol 98, 2023–2026, doi:10.1111/j.1572-0241.2003.07632.x (2003).

15 Hallert, C. et al. Evidence of poor vitamin status in coeliac patients on a gluten-free diet for 10 years. Aliment Pharmacol Ther 16, 1333–1339, doi:10.1046/j.1365-2036.2002.01283.x (2002).

16 Gensollen, T., Iyer, S. S., Kasper, D. L. & Blumberg, R. S. How colonization by microbiota in early life shapes the immune system. Science 352, 539–544, doi:10.1126/science.aad9378 (2016).

17 Atarashi, K. & Honda, K. Microbiota in autoimmunity and tolerance. Curr Opin Immunol 23, 761–768, doi:10.1016/j.coi.2011.11.002 (2011).

18 Getts, D. R., Chastain, E. M., Terry, R. L. & Miller, S. D. Virus infection, antiviral immunity, and autoimmunity. Immunol Rev 255, 197–209, doi:10.1111/imr.12091 (2013).

19 Vojdani, A. A Potential Link between Environmental Triggers and Autoimmunity. Autoimmune Dis 2014, 437231, doi:10.1155/2014/437231 (2014).

20 Nistal, E. et al. Differences in faecal bacteria populations and faecal bacteria metabolism in healthy adults and celiac disease patients. Biochimie 94, 1724–1729, doi:10.1016/j.biochi.2012.03.025 (2012).

21 Valitutti, F., Cucchiara, S. & Fasano, A. Celiac Disease and the Microbiome. Nutrients 11, doi:10.3390/nu11102403 (2019).

22 Zafeiropoulou, K. et al. Alterations in Intestinal Microbiota of Children With Celiac Disease at the Time of Diagnosis and on a Gluten-free Diet. Gastroenterology 159, 2039–2051 e2020, doi:10.1053/j.gastro.2020.08.007 (2020).

23 Cheng, J. et al. Duodenal microbiota composition and mucosal homeostasis in pediatric celiac disease. BMC Gastroenterol 13, 113, doi:10.1186/1471-230X-13-113 (2013).

24 Collado, M. C., Donat, E., Ribes-Koninckx, C., Calabuig, M. & Sanz, Y. Specific duodenal and faecal bacterial groups associated with paediatric coeliac disease. J Clin Pathol 62, 264–269, doi:10.1136/jcp.2008.061366 (2009).

25 De Palma, G. et al. Intestinal dysbiosis and reduced immunoglobulin-coated bacteria associated with coeliac disease in children. BMC Microbiol 10, 63, doi:10.1186/1471-2180-10-63 (2010).

26 Sanz, Y. et al. Differences in faecal bacterial communities in coeliac and healthy children as detected by PCR and denaturing gradient gel electrophoresis. FEMS Immunol Med Microbiol 51, 562–568, doi:10.1111/j.1574-695X.2007.00337.x (2007).

27 Olivares, M. et al. Gut microbiota trajectory in early life may predict development of celiac disease. Microbiome 6, 36, doi:10.1186/s40168-018-0415-6 (2018).

28 Serena, G. et al. Proinflammatory cytokine interferon-gamma and microbiome-derived metabolites dictate epigenetic switch between forkhead box protein 3 isoforms in coeliac disease. Clin Exp Immunol 187, 490–506, doi:10.1111/cei.12911 (2017).

29 Sellitto, M. et al. Proof of concept of microbiome-metabolome analysis and delayed gluten exposure on celiac disease autoimmunity in genetically at-risk infants. PLoS One 7, e33387, doi:10.1371/journal.pone.0033387 (2012).

30 Leonard, M. M. et al. Microbiome signatures of progression toward celiac disease onset in at-risk children in a longitudinal prospective cohort study. Proc Natl Acad Sci U S A 118, doi:10.1073/pnas.2020322118 (2021).

31 Amr, K. S., Bayoumi, F. S., Eissa, E. & Abu-Zekry, M. Circulating microRNAs as potential non-invasive biomarkers in pediatric patients with celiac disease. Eur Ann Allergy Clin Immunol 51, 159–164, doi:10.23822/EurAnnACI.1764-1489.90 (2019).

32 Sanchez, E., Donat, E., Ribes-Koninckx, C., Calabuig, M. & Sanz, Y. Intestinal Bacteroides species associated with coeliac disease. J Clin Pathol 63, 1105–1111, doi:10.1136/jcp.2010.076950 (2010).

33 Schippa, S. et al. A distinctive ’microbial signature’ in celiac pediatric patients. BMC Microbiol 10, 175, doi:10.1186/1471-2180-10-175 (2010).

34 Girdhar, K. et al. Dynamics of the gut microbiome, IgA response, and plasma metabolome in the development of pediatric celiac disease. Microbiome 11, 9, doi:10.1186/s40168-022-01429-2 (2023).

35 Cole, J. R. et al. Ribosomal Database Project: data and tools for high throughput rRNA analysis. Nucleic Acids Res 42, D633–642, doi:10.1093/nar/gkt1244 (2014).

36 McMurdie, P. J. & Holmes, S. phyloseq: an R package for reproducible interactive analysis and graphics of microbiome census data. PLoS One 8, e61217, doi:10.1371/journal.pone.0061217 (2013).

37 Oje, a. vegan: community ecology package. R’ ’ package. The Comprehensive R Archive Network (CRAN) (2019).

38 Robinson, M. D., McCarthy, D. J. & Smyth, G. K. edgeR: a Bioconductor package for differential expression analysis of digital gene expression data. Bioinformatics 26, 139–140, doi:10.1093/bioinformatics/btp616 (2010).

39 Yh, B. Controlling the false discovery rate: A practical and powerful approach to multiple testing. Journal of the Royal Statistical Society: Series B (Methodological) 57, 289–300 (1995).

40 Ye, Y. & Doak, T. G. A parsimony approach to biological pathway reconstruction/inference for genomes and metagenomes. PLoS Comput Biol 5, e1000465, doi:10.1371/journal.pcbi.1000465 (2009).

41 Chow, Y. W., Pietranico, R. & Mukerji, A. Studies of oxygen binding energy to hemoglobin molecule. Biochem Biophys Res Commun 66, 1424–1431, doi:10.1016/0006-291x(75)90518-5 (1975).

42 Douglas, G. PICRUSt2: An improved and customizable approach for metagenome inference. *BioRxiv* (2020).

43 Ritchie, M. E. et al. limma powers differential expression analyses for RNA-sequencing and microarray studies. Nucleic Acids Research 43, e47–e47, doi:10.1093/nar/gkv007 (2015).

44 WIckham, H. ggplot2 - Elegant Graphics for Data Analysis. Springer International Publishing doi:10.1101/672295 (2016).

45 RK, K. pheatmap: Implementation of heatmaps that offers more control over dimensions and appearance. The Comprehensive R Archive Network (CRAN) (2019).

46 Koenig, J. E. et al. Succession of microbial consortia in the developing infant gut microbiome. Proc Natl Acad Sci U S A 108 **Suppl 1**, 4578–4585, doi:10.1073/pnas.1000081107 (2011).

47 Langille, M. G. et al. Predictive functional profiling of microbial communities using 16S rRNA marker gene sequences. Nat Biotechnol 31, 814–821, doi:10.1038/nbt.2676 (2013).

48 Tan, F. Y., Tang, C. M. & Exley, R. M. Sugar coating: bacterial protein glycosylation and host-microbe interactions. Trends Biochem Sci 40, 342–350, doi:10.1016/j.tibs.2015.03.016 (2015).

49 Matilla, M. A. & Krell, T. The effect of bacterial chemotaxis on host infection and pathogenicity. FEMS Microbiol Rev 42, doi:10.1093/femsre/fux052 (2018).

50 Aharonowitz, Y., Cohen, G. & Martin, J. F. Penicillin and cephalosporin biosynthetic genes: structure, organization, regulation, and evolution. Annu Rev Microbiol 46, 461–495, doi:10.1146/annurev.mi.46.100192.002333 (1992).

51 Moore, A. M. et al. Gut resistome development in healthy twin pairs in the first year of life. Microbiome 3, 27, doi:10.1186/s40168-015-0090-9 (2015).

52 Bakhtiari, S. et al. The connection between fatty acids and inflammation in celiac disease; a deep exploring. Tissue Barriers, 2342619, doi:10.1080/21688370.2024.2342619 (2024).

53 Solakivi, T. et al. Serum fatty acid profile in celiac disease patients before and after a gluten-free diet. Scand J Gastroenterol 44, 826–830, doi:10.1080/00365520902912589 (2009).

54 Riezzo, G. et al. Lipidomic analysis of fatty acids in erythrocytes of coeliac patients before and after a gluten-free diet intervention: a comparison with healthy subjects. Br J Nutr 112, 1787–1796, doi:10.1017/S0007114514002815 (2014).

55 Khalkhal, E. et al. Screening of Altered Metabolites and Metabolic Pathways in Celiac Disease Using NMR Spectroscopy. Biomed Res Int 2021, 1798783, doi:10.1155/2021/1798783 (2021).

56 Girdhar, K. et al. A gut microbial peptide and molecular mimicry in the pathogenesis of type 1 diabetes. Proc Natl Acad Sci U S A 119, e2120028119, doi:10.1073/pnas.2120028119 (2022).

57 Dedrick, S. et al. The Role of Gut Microbiota and Environmental Factors in Type 1 Diabetes Pathogenesis. Front Endocrinol (Lausanne) 11, 78, doi:10.3389/fendo.2020.00078 (2020).

58 Marietta, E., Horwath, I., Balakrishnan, B. & Taneja, V. Role of the intestinal microbiome in autoimmune diseases and its use in treatments. Cell Immunol 339, 50–58, doi:10.1016/j.cellimm.2018.10.005 (2019).

59 Ordonez-Rodriguez, A., Roman, P., Rueda-Ruzafa, L., Campos-Rios, A. & Cardona, D. Changes in Gut Microbiota and Multiple Sclerosis: A Systematic Review. Int J Environ Res Public Health 20, doi:10.3390/ijerph20054624 (2023).

60 Christovich, A. & Luo, X. M. Gut Microbiota, Leaky Gut, and Autoimmune Diseases. Front Immunol 13, 946248, doi:10.3389/fimmu.2022.946248 (2022).

61 De Luca, F. & Shoenfeld, Y. The microbiome in autoimmune diseases. Clin Exp Immunol 195, 74–85, doi:10.1111/cei.13158 (2019).

62 D’Argenio, V. et al. Metagenomics Reveals Dysbiosis and a Potentially Pathogenic N. flavescens Strain in Duodenum of Adult Celiac Patients. Am J Gastroenterol 111, 879–890, doi:10.1038/ajg.2016.95 (2016).

63 Wacklin, P. et al. The duodenal microbiota composition of adult celiac disease patients is associated with the clinical manifestation of the disease. Inflamm Bowel Dis 19, 934–941, doi:10.1097/MIB.0b013e31828029a9 (2013).

64 Stewart, C. J. et al. Temporal development of the gut microbiome in early childhood from the TEDDY study. Nature 562, 583–588, doi:10.1038/s41586-018-0617-x (2018).

65 Lionetti, E. et al. Introduction of gluten, HLA status, and the risk of celiac disease in children. N Engl J Med 371, 1295–1303, doi:10.1056/NEJMoa1400697 (2014).

66 Caminero, A. et al. Diversity of the cultivable human gut microbiome involved in gluten metabolism: isolation of microorganisms with potential interest for coeliac disease. FEMS Microbiol Ecol 88, 309–319, doi:10.1111/1574-6941.12295 (2014).

67 Sample, D. et al. Baseline Fecal Microbiota in Pediatric Patients With Celiac Disease Is Similar to Controls But Dissimilar After 1 Year on the Gluten-Free Diet. JPGN Rep 2, e127, doi:10.1097/PG9.0000000000000127 (2021).

68 Leonard, M. M. et al. Multi-omics analysis reveals the influence of genetic and environmental risk factors on developing gut microbiota in infants at risk of celiac disease. Microbiome 8, 130, doi:10.1186/s40168-020-00906-w (2020).

69 Singh, P. et al. Distinctive Microbial Signatures and Gut-Brain Crosstalk in Pediatric Patients with Coeliac Disease and Type 1 Diabetes Mellitus. Int J Mol Sci 22, doi:10.3390/ijms22041511 (2021).

70 Bonder, M. J. et al. The influence of a short-term gluten-free diet on the human gut microbiome. Genome Med 8, 45, doi:10.1186/s13073-016-0295-y (2016).

71 Olivares, M. et al. The HLA-DQ2 genotype selects for early intestinal microbiota composition in infants at high risk of developing coeliac disease. Gut 64, 406–417, doi:10.1136/gutjnl-2014-306931 (2015).

72 Milletich, P. L. et al. Gut microbiome markers in subgroups of HLA class II genotyped infants signal future celiac disease in the general population: ABIS study. Front Cell Infect Microbiol 12, 920735, doi:10.3389/fcimb.2022.920735 (2022).

73 Collado, M. C., Donat, E., Ribes-Koninckx, C., Calabuig, M. & Sanz, Y. Imbalances in faecal and duodenal Bifidobacterium species composition in active and non-active coeliac disease. BMC Microbiol 8, 232, doi:10.1186/1471-2180-8-232 (2008).

74 Girdhar, K. et al. Dynamics of Gut Microbiome, IgA Response and Plasma Metabolome in Development of Pediatric Celiac Disease. bioRxiv, 2020.2002.2029.971242, doi:10.1101/2020.02.29.971242 (2022).

75 Chow, M. A., Lebwohl, B., Reilly, N. R. & Green, P. H. Immunoglobulin A deficiency in celiac disease. J Clin Gastroenterol 46, 850–854, doi:10.1097/MCG.0b013e31824b2277 (2012).

76 Planer, J. D. et al. Development of the gut microbiota and mucosal IgA responses in twins and gnotobiotic mice. Nature 534, 263–266, doi:10.1038/nature17940 (2016).

77 Stojiljkovic, V. et al. Glutathione redox cycle in small intestinal mucosa and peripheral blood of pediatric celiac disease patients. An Acad Bras Cienc 84, 175–184, doi:10.1590/s0001-37652012000100018 (2012).

78 Pavlick, K. P. et al. Role of reactive metabolites of oxygen and nitrogen in inflammatory bowel disease. Free Radic Biol Med 33, 311–322, doi:10.1016/s0891-5849(02)00853-5 (2002).

79 Leclerc, M. et al. Nitric Oxide Impacts Human Gut Microbiota Diversity and Functionalities. mSystems 6, e0055821, doi:10.1128/mSystems.00558-21 (2021).

80 Baldi, S. et al. Free Fatty Acids Signature in Human Intestinal Disorders: Significant Association between Butyric Acid and Celiac Disease. Nutrients 13, doi:10.3390/nu13030742 (2021).

81 de Jong, A. J., Kloppenburg, M., Toes, R. E. & Ioan-Facsinay, A. Fatty acids, lipid mediators, and T-cell function. Front Immunol 5, 483, doi:10.3389/fimmu.2014.00483 (2014).

82 Tjellstrom, B. et al. Faecal short-chain fatty acid pattern in childhood coeliac disease is normalised after more than one year’s gluten-free diet. Microb Ecol Health Dis 24, doi:10.3402/mehd.v24i0.20905 (2013).

83 Mucida, D. et al. Reciprocal TH17 and regulatory T cell differentiation mediated by retinoic acid. Science 317, 256–260, doi:10.1126/science.1145697 (2007).

84 Fallah, S. et al. Investigating the Impact of Vitamin A and Amino Acids on Immune Responses in Celiac Disease Patients. Diseases 12, doi:10.3390/diseases12010013 (2024).

85 DePaolo, R. W. et al. Co-adjuvant effects of retinoic acid and IL-15 induce inflammatory immunity to dietary antigens. Nature 471, 220–224, doi:10.1038/nature09849 (2011).

86 Winslet, M. C., Poxon, V., Allan, A. & Keighley, M. R. Mucosal glucosamine synthetase activity in inflammatory bowel disease. Dig Dis Sci 39, 540–544, doi:10.1007/bf02088339 (1994).

87 Mayassi, T. et al. Chronic Inflammation Permanently Reshapes Tissue-Resident Immunity in Celiac Disease. Cell 176, 967–981 e919, doi:10.1016/j.cell.2018.12.039 (2019).

